# Genome-wide analysis of H3K4me3 and H3K27me3 modifications throughout the mouse urogenital ridge at E11.5

**DOI:** 10.1101/347880

**Authors:** Yisheng Yang, Megan J Wilson

## Abstract

In mammals, the adrenal gland, testis and ovary arise from a common progenitor tissue known as the urogenital ridge (UGR). This small population of cells will adopt a number of different cell fates following sex determination, including forming the precursors of somatic cells (such as Sertoli and granulosa cells) and steroidogenic cells. In addition, this tissues also contains the Wolffian and Müllerian ducts that later form components of the reproductive tracts. A potential mechanism to maintain developmental plasticity of the UGR until gonad formation is through the epigenetic modification of histone proteins.

In order to provide a resource for future studies, we used chromatin immunoprecipitation followed by high throughput sequencing (ChIP-seq) for two histone modifications, H3K4me3 and H3K27me3, in the E11.5 mouse UGR. These marks are both known to reflect the active, repressive or a poised chromatin state. We found that enrichment for each histone mark reflected transcriptional activity in precursor cells of the developing gonad. From the analysis of potential enhancer/regulator peak regions for DNA binding motifs, we identified several candidate transcription factors that may contribute to gonadal cell lineage specification. We additionally identified signaling pathway genes that are targeted by both chromatin modifications. Together, these datasets provide a useful resource for investigating gene regulatory networks functioning during UGR development at E11.5.

## Introduction

The mammalian urogenital ridge (UGR) is a unique developmental structure that has the ability to adopt two quite distinct fates during late embryonic development. Each bi-potential UGR consists of the progenitor cells required to form either a testis or ovary: the supporting cells (Sertoli or granulosa cells), the germ cells (GCs), the interstitial cells (including steroidogenic cells) and endothelial cells (Capel, 2017). Cell lineage RNA analysis (Jameson et al., 2012) has suggested that the gonadal progenitor cells are biased towards one of two cell fates: germ cells are considered male-biased, whereas supporting cells are female-biased. The supporting cells adopt a sex-specific fate first, and this process is completed by embryonic day 12.5 (E12.5) in mice. The germ and interstitial cells follow, starting at E12.5 (Jameson et al., 2012). How cell fate plasticity is maintained within the developing UGR prior to and during early sex determination is unknown.

Sex determination in mammals is a critical step in specification of gonadal cell fate and depends on the chromosomal makeup of the embryo. The presence of a Y chromosome (XY genotype) shifts the bi-potential UGR towards a testicular fate, through the initial expression of the testis determining gene, *Sex determining region Y* (*Sry*). *Sry* in turn activates *Sry-box 9* (*Sox9*) gene expression required for Sertoli cell differentiation and the activation of signaling pathways for the maturation of testis-specific cell types (Sekido and Lovell-Badge, 2008; Wilhelm et al., 2007; Wilson et al., 2005). SOX9 up-regulates *Anti-Müllerian hormone* (*AMH*) to promote the regression of the Müllerian ducts, whereas the Wolffian ducts develop into male-specific structures under the influence of testosterone, produced by the Leydig cells (Munsterberg and Lovell-Badge, 1991; Wilhelm et al., 2006).

In the absence of a Y chromosome (XX genotype), the female developmental pathway is initiated. Unlike male development, the morphology of the gonad does not change drastically in the immediate period after initiation of the female pathway (Mork et al., 2012). Pre-granulosa cell specification occurs at ~E12.5 accompanied by expression of *Forkhead box L2* (*Foxl2*) (Schmidt et al., 2004). *Sox9* gene expression is repressed in the XX gonad by ovary-specific Wnt4/β-catenin/Rspo1 signaling pathway that promotes the expression of *Foxl2* and the differentiation of the Müllerian ducts into female-specific structures (Chassot et al., 2008; Pannetier et al., 2016; Tanaka and Nishinakamura, 2014). The primordial GCs arise near the yolk sac and migrate into the UGR, arriving at ~E10.5 (Molyneaux et al., 2001). GCs located in the developing ovary enter meiosis due to the presence of retinoic acid (RA) produced from the neighboring mesonephros. In contrast, male germ cells are prevented from beginning meiosis, as RA is degraded in the developing testis by the CYP26B1 enzyme (Bowles et al., 2006; Spiller et al., 2017).

Due to the bi-potential nature of the UGR, in some disorders of sex development (DSD) an ovotestis may result, a mix of both male and female associated cell types located within a single adult gonad. This disorders are often a result of mutations in the *SRY* or *SOX9* genes (Vilain, 2011). Conditional knockout of *Foxl2* in the adult ovary results in the formation of Sertoli- and Leydig-like cells, suggesting a role for *Foxl2* in maintaining the ovarian phenotype in the adult as well as the embryo (Uhlenhaut et al., 2009). Likewise, inactivation of *DMRT1* in adult Sertoli cells increases *Foxl2* gene expression, producing granulosa and theca-like cells, along with the production of estrogen (Matson et al., 2011). Together, these studies demonstrate the incredible plasticity of sex-development and the need to continually maintain one pathway (either male or female) over the other throughout the adult lifetime.

One possible mechanism to regulate and maintain gonadal cell fate is through the use of specific histone modifications. Global genome-wide changes to histone modifications could also be involved in initiating differentiation of precursor cells within the gonad and mesonephros. Epigenetic modifications play key roles in cell lineage specification during development by manipulating chromatin structure and altering gene expression (Atlasi and Stunnenberg, 2017). Chromatin is found in two states: euchromatin, an open, unwound formation that allows for an active state of transcription, and heterochromatin, a closed, tightly wound formation that represses transcription (Parker et al., 2004). Post-translational modifications to histone proteins largely occur to amino acids located at the N-terminal tail, influencing DNA accessibility to transcriptional regulators of gene expression. Trimethylation of lysine N-terminal residues can induce either an active or repressed chromatin configuration depending on location. Histone 3 lysine 4 trimethylation (H3K4me3) is a hallmark of actively transcribed genes and is commonly associated with transcription start sites (TSS) and promoter regions, whereas histone 3 lysine 27 trimethylation (H3K27me3) is strongly associated with inactive promoter regions and repressed gene transcription (Bannister and Kouzarides, 2011). Both active and repressive marks can be present at the same genomic regions, indicating a poised state common in undifferentiated cells such as embryonic stem cells (Bernstein et al., 2006; Voigt et al., 2013).

There is limited data regarding the epigenetic profile of gonadal somatic and germ cells *in vivo*. Using cell sorting, researchers have isolated the primordial GCs (PGCs) using Oct4-GFP mouse line (Ng et al., 2013; Sachs et al., 2013) to study epigenetic modifications using chromatin immunoprecipitation followed by sequencing (ChIP-seq). PGCs tend to have high levels of the repressive H3K27me3 mark and are transcriptionally silent (Ng et al., 2013; Sachs et al., 2013). Bivalent regions, areas with both active and repressive histone modifications, are enriched within PGCs and tend to be located near developmental genes (Sachs et al., 2013). Recently, DNaseI-seq and ChIP-seq for H3K27ac was used to identify active enhancer elements in Sertoli cells at E15.5 (Maatouk et al., 2017).

To obtain a more global view of histone modifications in all cells of the developing bi-potential gonad, we used ChIP-seq to study the distribution of H3K27me3 and H3K4me3 in E11.5 gonads. Profiling of histone mortifications not only gives information about the transcriptional activity of nearby genes, but can aid in the identification of tissue-specific enhancer elements. Through ChIP-seq, we discovered different histone signal profiles that reflect gene expression levels in the E11.5 gonad. Using this data, we identified overrepresented DNA binding motifs to determine candidate transcription factors, and then extracted their cell-type expression values during gonad development using publically available data (Jameson et al., 2012). This data set provides a resource for future studies of early gonadal development.

## Results

### Immunostaining of UGR at E11.5

Sectioned E11.5 embryos were stained for H3K4me3 and H3K27me3 histone marks (Fig. 1). While all cells were positive for some level of these histone modifications, there were many cells located within the gonad that stained much more strongly than nearby cells (Fig. 1). These strongly-staining cells were located along the coelomic epithelium and scattered throughout the gonad (Fig. 1A). While some of these cells are likely to be primordial GCs, many of these cells are also somatic cells. For instance, cells that line the coelomic epithelium are precursors of supporting cell lineages such as Sertoli cells (Karl and Capel, 1998). Staining for each histone modification was found in the nucleus, often co-localized at nuclear foci (Fig. 1B).

**Figure 1.**
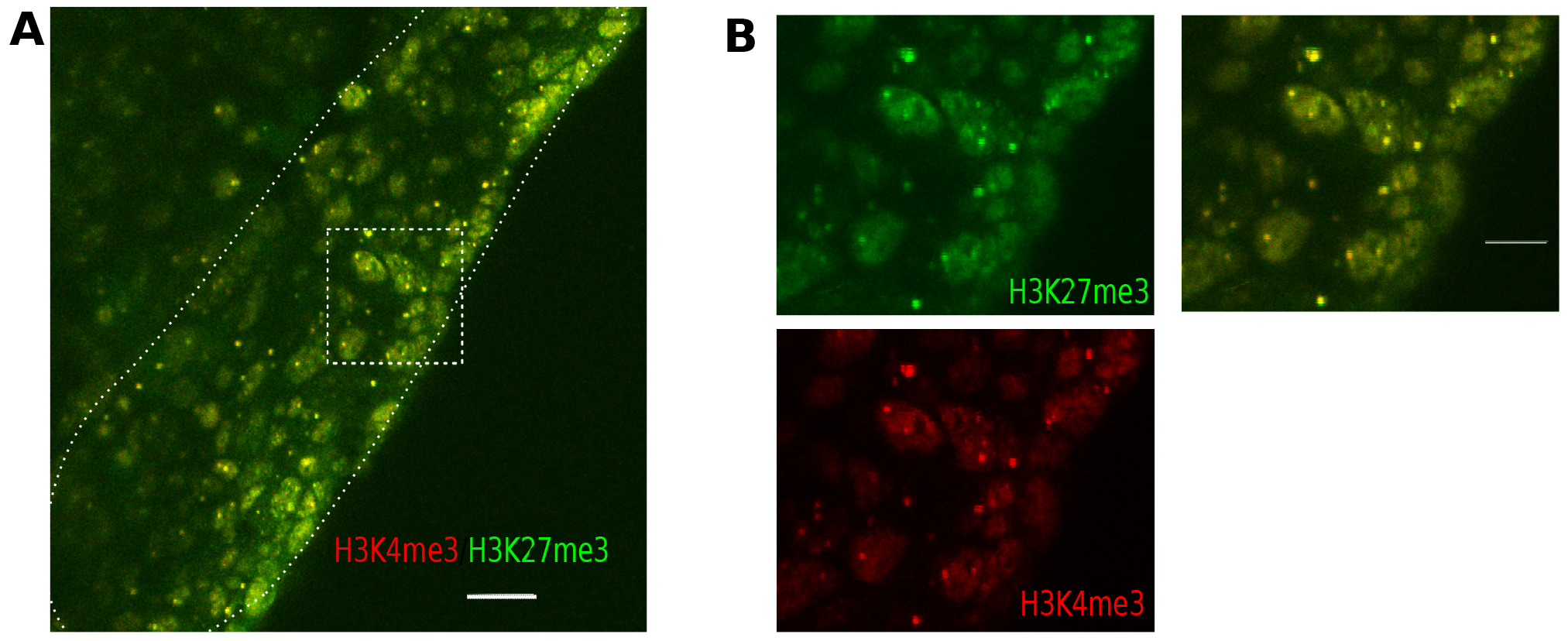
Detection of histone marks H3K27me3 and H3K4me3 marks in the UGR. **A.** Double immunofluoresence staining for H3K27me3 (green) and H3K4me3 (red). The gonad is outlined with a dotted line. Scale bar = 20 μm. Dashed box indicates region shown in **B**.

### Genome-wide identification of H3K27me3 and H3K4me3

Urogenital ridge tissue was dissected from each E11.5 stage embryo and pooled (two biological replicates, five litters per replicate,), prior to cross-linking briefly with formaldehyde. Extracted chromatin was fragmented by MNase digestion, followed by immunoprecipitation using antibodies for either H3K27me3 or H3K4me3. Two biological replicates (from separate ChIP experiments) were sequenced on an Illumina platform to detect each histone mark (our approach is summarized in Fig. 2A). Sequencing reads were aligned to the mouse reference genome (mm9, File 1) and correlation analysis revealed good reproducibility between each biological replicate (>93% Pearson correlation, Fig. S1). For global signal distribution analysis, both replicates were pooled.

**Figure 2.**
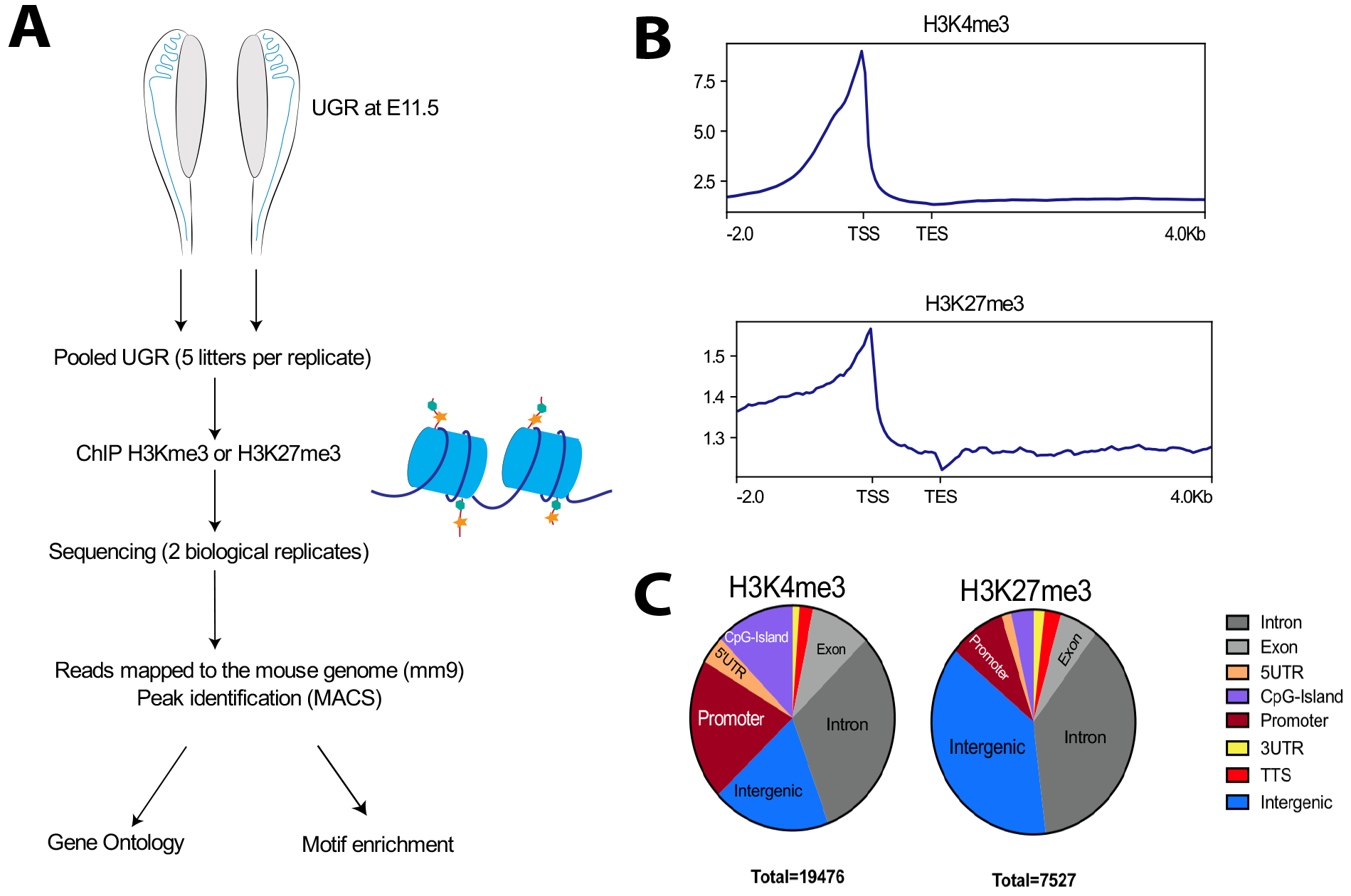
ChIP-seq for H3K4me3 and H4K27me3 histone marks. **A.** Overview of the ChIP-sequencing experiments. **B.** Signal profile with respect to the average gene body and 2 kb upstream, 4kb downstream. **C.** ChIP-seq peak location with respect to genome annotations of protein coding genes. Abbreviations: Transcriptional start site (TSS), transcription end site (TES).

We used MACS broad peak caller (bdgbroadcall) to call peak regions, genomic regions with statistically enriched signal compared to the input controls. This identified 32,182 peak regions for H3K4me3 (with an average peak length of 1809 bp) and 13,556 regions for H3K27me3 (with an average length of 1310 bp). Peak location, with respect to gene body features, was analyzed with HOMER (Heinz et al., 2010) and CEAS (Shin et al., 2009) for each histone mark. Of the MACS broadcall regions, 19476 (H3K4me3) and 7527 (H3K27me3) peaks were located near known RefSeq genes. H3K4me3 was more enriched at the 5’ end of genes, near the transcriptional start site (TSS) (Fig. 2B and Fig. S2) including CpG islands (Fig 2C). The repressive histone mark, H3K27me3 was more generally enriched for within intergenic regions and introns, and a smaller proportion of peaks were located within promoter regions (Fig. 2B and 2C, Fig. S2). This data is consistent with previous studies, showing that H3K4me3 is found at active promoter regions and enhancers (Santos-Rosa et al., 2002), whereas the H3K27me3 mark is largely distributed across the bodies of genes, with some limited enrichment near the TSS, often at bivalently marked genes (Young et al., 2011).

The distribution of histone marks at five key sex-determining genes was examined further (Fig 3. and Fig. S3). Active histone modification enrichment near promoter regions, and within either an intron or nearby intergenic regions, can indicate the presence of an enhancer region (Atlasi and Stunnenberg, 2017). *Sox9*, *Wt1* and *Sf1* genes are all expressed in the developing UGR at E11.5 and, following sex-determination, play important roles in testicular development (Hammes et al., 2001; Kent et al., 1996; Luo et al., 1994). *Lhx9* and *Cbx2* are required for formation of the mouse gonad primordium (Birk et al., 2000; Katoh-Fukui et al., 2005). The *Sf1* (also known as *Nr5a1*) gene showed enrichment for both histone marks, peak regions were located within the 4^th^ intron and near the promoter region of the gene (Fig. 3A). H3K4me3 signal was strongly enriched for around the promoter region and first exons for *Wt1* and *Wt1os* genes (Fig. 3B). *Wt1os* encodes a long non-coding RNA, co-expressed with *Wt1* in many tissues (Dallosso et al., 2007). A narrow peak region for H3K27me3 signal, overlapping with H3K3me4 signal, was also identified in the 2^nd^ intron (Fig. 3B). An additional H3K4me3 peak region is located ~3 kb, 3’ of the *Wt1* gene (Fig. 3B); this may represent an active enhancer element.

**Figure 3.**
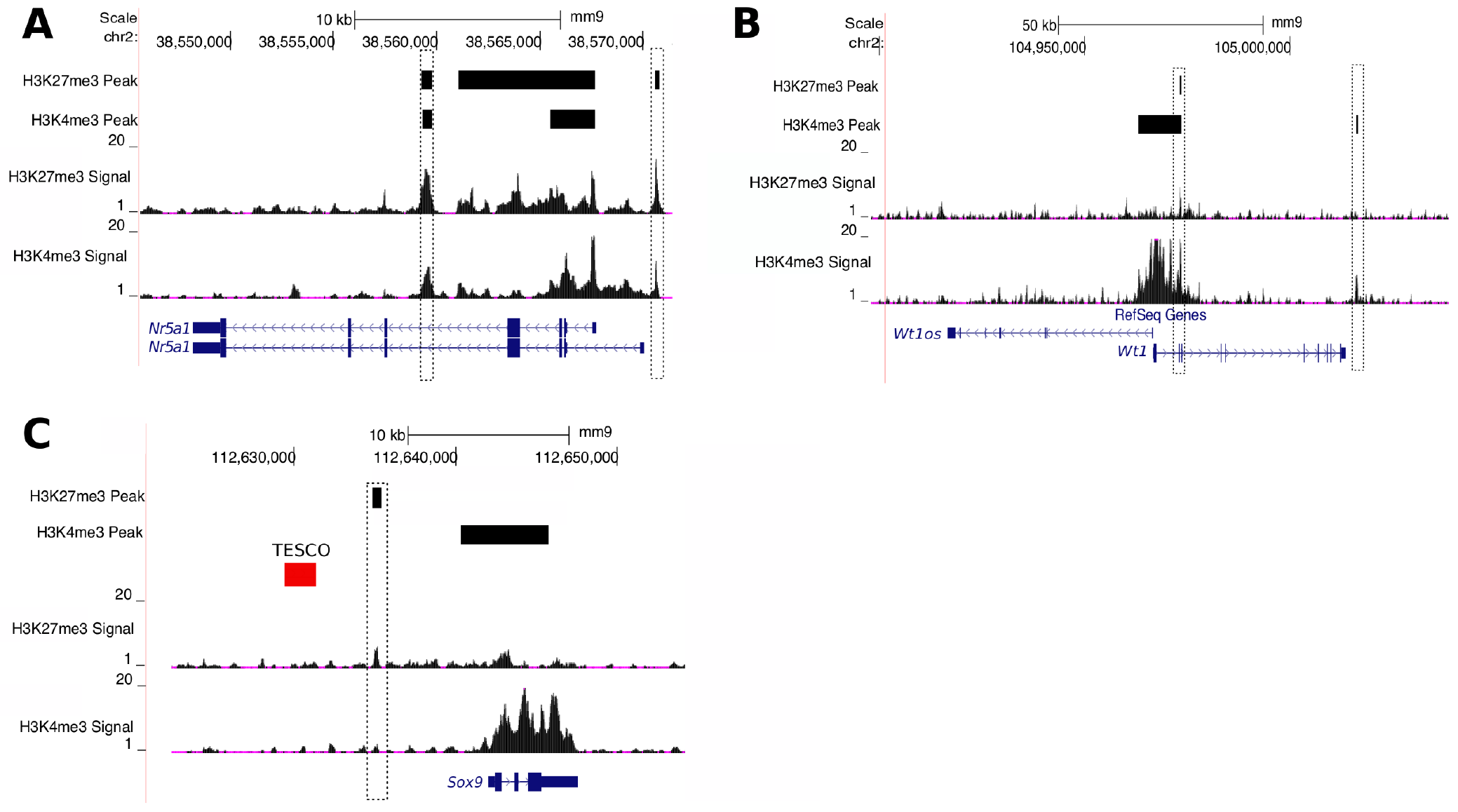
Examples of signal distribution near three genes required for early gonad formation and sex-determination. **A.** *SF1/Nr5a1* gene **B.** *Wt1* gene. **C.** *Sox9* gene. The TESCO enhancer (red bar) is also shown for the *Sox9* gene. Signal tracks for each histone mark are shown above each gene model. Black bars indicate peak regions identified by MACS analysis. Boxed region indicates enriched signal for both histone marks and therefore, the location of a putative enhancer.

*Sox9* is a key male sex-determination gene and target of *Sry* (Kent et al., 1996; Sekido and Lovell-Badge, 2008). The H3K4me3 modification was strongly enriched around the *Sox9* gene (Fig. 3C), reflecting the fact that this gene is expressed in both sexes at E11.5 (Kent et al., 1996). There was an additional peak for H3K27me3, located 8.4 kb upstream of the *Sox9* promoter. This genomic region is ~3.5 kbp away from the TESCO (testis-specific enhancer of *Sox9* core element) enhancer, thought to restrict *Sox9* expression to the testis (Bernard et al., 2012; Gonen et al., 2017; Sekido and Lovell-Badge, 2008).

*Lhx9* and *Cbx2* (also known as *M33*) genes are required for formation of the UGR and are expressed at E11.5 (Birk et al., 2000; Katoh-Fukui et al., 2005). H3K4me3 signal was high around the TSS for both genes (Fig. S3), indicating active promoter regions. There was also a peak for *2310009B15Rik*, a protein-coding gene of unknown function located near the *Lhx9* gene, suggesting that this gene is also expressed at E11.5. Signal for the repressive histone mark H3K27me3 was also found near the Lhx9 TSS and throughout the gene body (Fig. S3A), indicating that in some gonadal cells *Lhx9* is repressed. Previous spatial expression studies have shown that *Lhx9* mRNA is strongest in the cells of the coelomic epithelium, and absent from some of the cells of the inner mesenchyme and mesonephros (Birk et al., 2000). In contrast, H3K27me3 ChIP-seq signal was low around the *Cbx2* gene, with no peak regions being identified by MACS (Fig. S3B), indicating little repression by H3K27me3 modification in the UGR cell population. *Cbx2* exhibits broader cell expression, being found in all cells of the gonad and the neighboring mesonesphros (Katoh-Fukui et al., 2012).

### Levels of H3K4me3 and H3K27me3 modifications reflect gene expression levels in the UGR

To identify genes with similar histone mark profiles, heatmaps with K-means clustering were generated using deepTools (Ramirez et al., 2014) to visualize the region around the TSS for RefSeq gene annotations and separate them into clusters with similar signal profiles (Fig. 4A and 5A, File 2). Gene ontology (GO) analysis was carried out for clustered genes using PANTHER to identify overrepresented biological annotations (Fig. 4B and 4C, Fig. 5B and 5C, Files 3 and 4). The expression levels of genes grouped within each signal profile cluster was also extracted from Affymetrix expression data previously generated for each cell lineage present in the E11.5 gonad by Jameson *et al.* (Jameson et al., 2012). This enabled us to determine the relationship between signal clusters and gene expression levels (Fig. 6).

**Figure 4.**
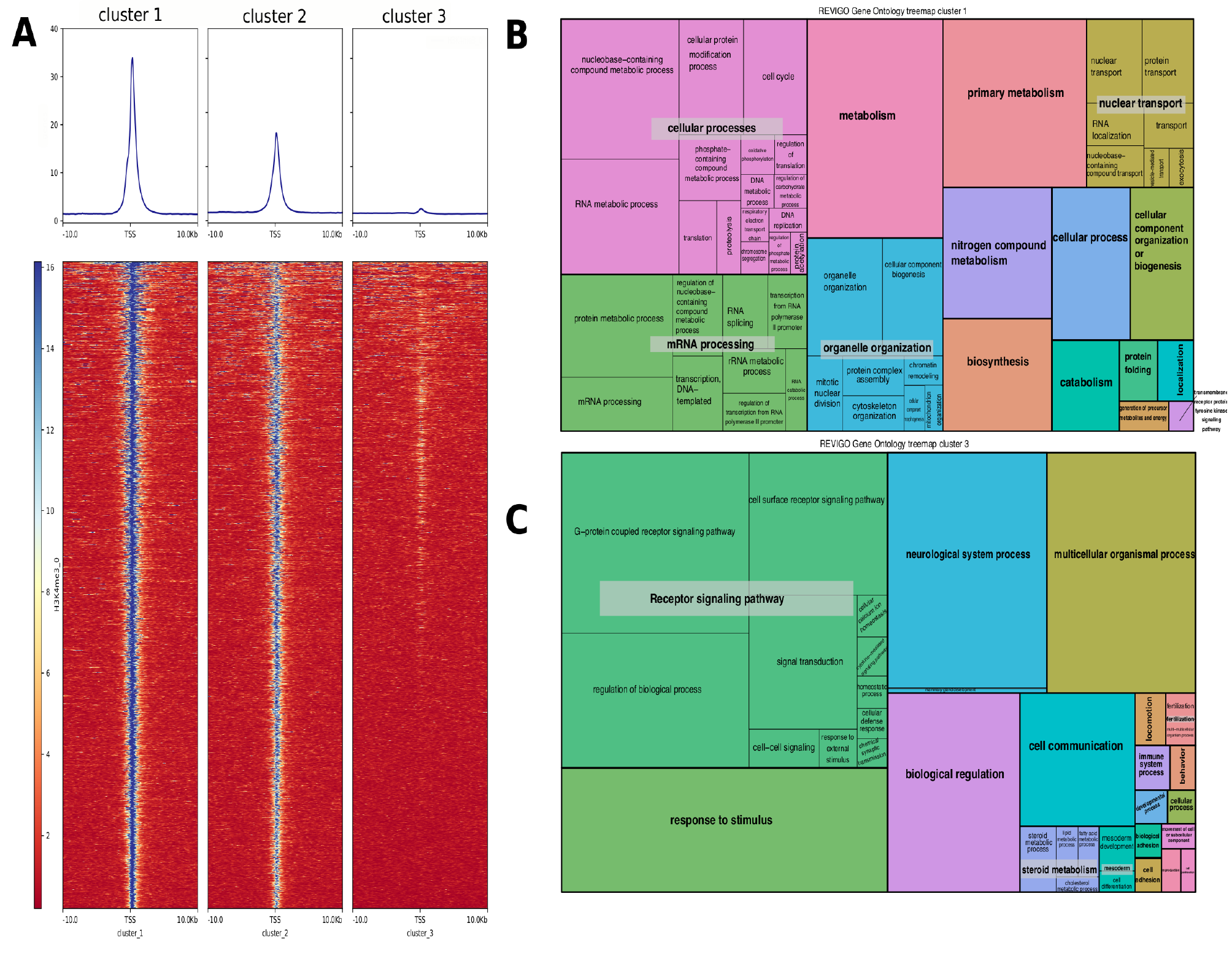
Heatmap profile for H3K4me3 and GO analysis for RefSeq genes cluster. **A.** Heatmap of H3K4me3 signal density using K-means clustering. Average signal profile is shown above each heatmap. Read counts were considered around the TSS (+/− 10 kb). Blue indicates low signal and Red indicates an area of high signal. **B.** Treemap summarizing GO analysis for cluster 1 genes (PANTHER slim biological processes). **C.** Treemap visualizing cluster 2 GO analysis (PANTHER slim biological processes). Boxes with similar GO-terms are grouped together and are displayed with the same color. The box size represents the −log10 P-value of GO-term enrichment. Additional Treemap for cluster 3 is provided in supplementary Fig. S4.

**Figure 5.**
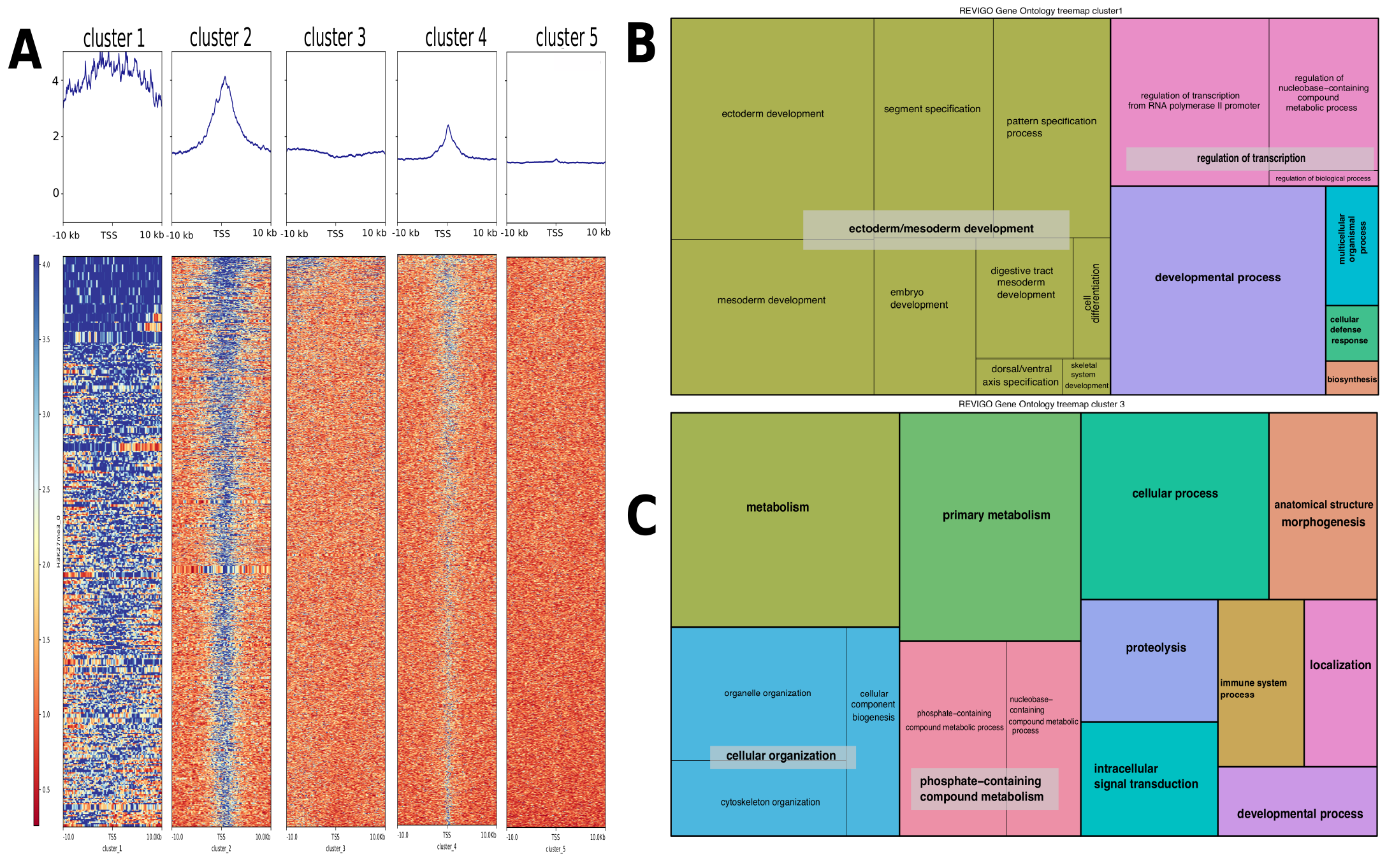
Cluster and GO analysis for H3K27me3 whole genome distribution. **A.** Signal distribution heatmap and K-means clustering for H3K27me3. Read counts were considered around the TSS (+/− 10 kb). Blue indicates low signal and Red indicates an area of high signal. **B.** Treemap of GO pathways (PANTHER slim biological processes) overrepresented for cluster 1. **C.** Treemap for cluster 3 enriched GO-terms (PANTHER slim biological processes). GO treemaps for the other signal clusters are provided in Fig. S5.

**Figure 6.**
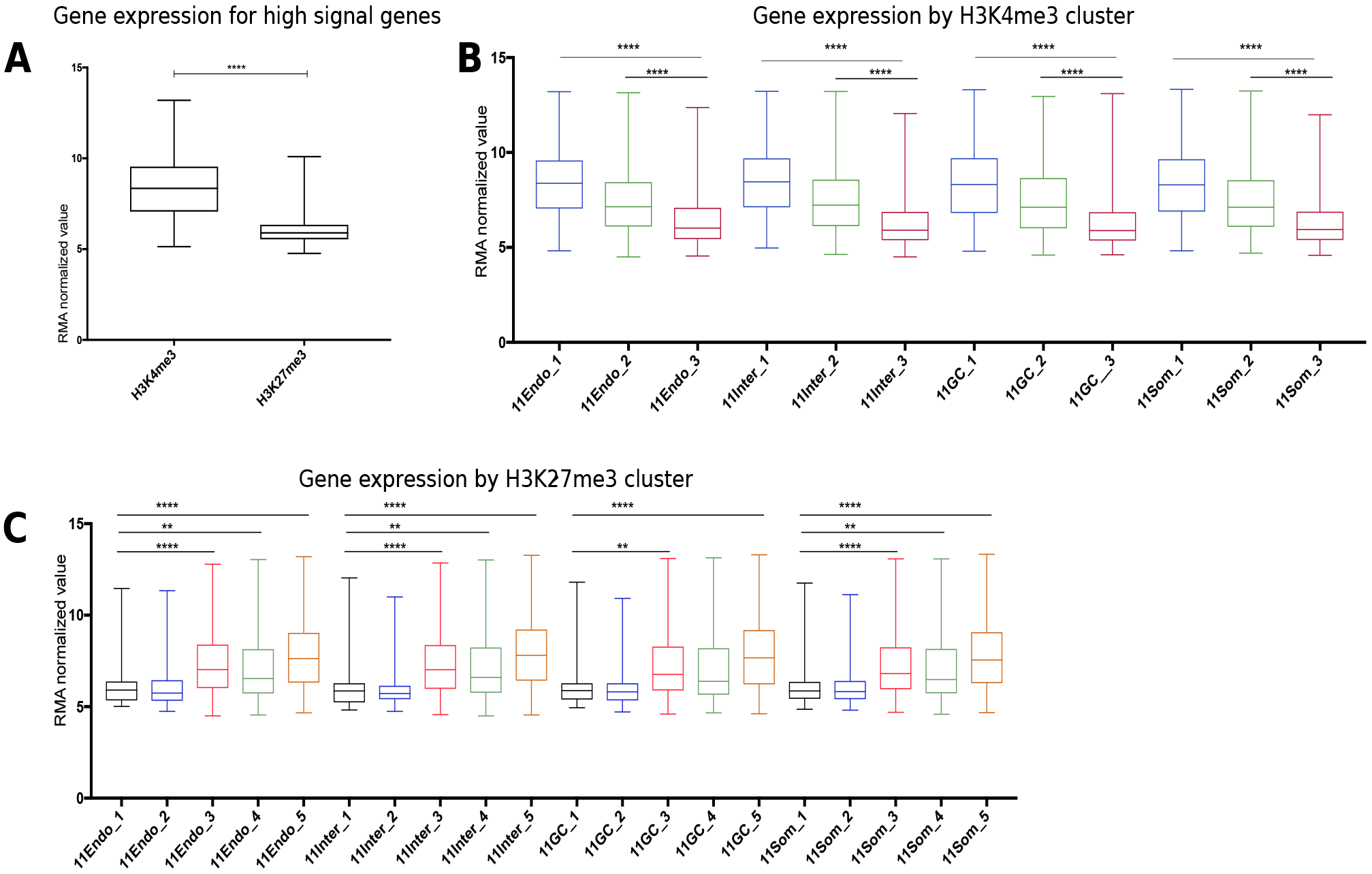
Gonadal expression of genes overlapping with distinct H3K4me3 and H3K27me3 signal profiles. **A.** Average gene expression for all cell types, for H4K4me3 cluster 1 and H3K27me3 cluster 2 genes. These two clusters have high signal density for each histone mark around the gene body. **B and C.** Expression of genes associated with each signal profile cluster, per cell lineage. One-way ANOVA Tukey’s multiple comparisons test adjusted P-value <0.001 (**), <0.0001 (****). Abbreviations: endothelial cells (Endo), germ cells (GC), interstitial/stroma (Inter), somatic cells (Som). Cluster number is indicated for each plot (eg 11Endo_1 is gene expression of cluster 1 genes in the endothelial cell lineage). Normalized gene expression data was taken from (Jameson et al., 2012).

Three clusters were generated for H3K4me3 signal distribution. The first cluster had high levels of H3K4me3 located near the gene body and TSS (Fig. 4A). Cluster 1 genes function in core cellular biological processes including general essential cellular processes such as RNA processing and metabolism (Fig. 4B). The second cluster (cluster 2) genomic regions have moderate levels of H3K4me3 signal (Fig. 4A), and are enriched for genes involved largely in sensory reception and receptor signaling genes (Fig. 4C). There are no significant differences in gene expression levels between cluster 1 and 2 genes (Fig. 6B). However, cluster 3 genes are expressed at significantly lower levels compared to cluster 1 and 2 genes (Fig. 5, two-way ANOVA, P < 0.0001). Genomic regions grouped into the third cluster have little low/no H3K4me3 signal (Fig. 4A). Out of 13,043 genes (18,043 RefSeq transcripts) associated within cluster 3 regions, only 3,392 genes are noted as expressed in the E11.5 gonad Affymetrix dataset (Jameson et al., 2012) (26% of genes, File 3). These genes function in a variety of different biological processes (Fig. S4A). In comparison, of the 6166 genes in cluster 1 (corresponding to 7005 RefSeq transcripts), 5056 of these are expressed in the E11.5 gonad (~82% of genes, File 3). This supports previous research showing that H3 genome-wide enrichment is linked to gene expression (Guenther et al., 2007; Mikkelsen et al., 2007).

H3K27me3 signal distribution can be divided into 5 clusters with different profiles. Cluster 1 histone mark covers over 20 kbp of DNA including the gene TSS (Fig. 5A), and is likely to represent areas of repressed chromatin. GO annotation revealed that these genome regions are enriched for genes that function in essential early developmental processes such as germ layer development and axis specification (Fig. 5B). Of these 219 cluster 1 genes, only 58 had detectable expression in single cell profiling (File 4), with expression levels much lower than genes found in other H3K27me3 signal profile clusters (Fig. 5A). The signal profile for cluster 2 genes is strongest around a +/− 5 kb region near the TSS. GO analysis revealed that many cluster 2 genes function in cell-cell signaling, differentiation and transcriptional regulation (Fig. 5C). This signal cluster is associated with 1559 genes (1750 RefSeq transcripts) and 33% of these genes (518 genes) are expressed in the gonad at E11.5 (File 4). As the H3K27me3 mark is associated with gene repression, we would expect expression of genes with high H3K27me3 signal, such as those genes associated with clusters 1 and 2, to be low or not detectable.

Cluster 3 had reduced levels of signal near the TSS compared to nearby regions (Fig. 5A, see enlarged view in Fig. S4B). These genes function in essential biological processes such as metabolism and sub-cellular organization (Fig 5C). Of 3844 genes with this signal profile, 56% (3138 genes) are expressed in the gonad at E11.5 (File 4). H3K27me3 signal profile for cluster 4 is confined to narrow region near the TSS (Fig. 5A). Cluster 4 associated genes are involved in the intracellular signaling including localization and transport of proteins (Fig. S5). Of the 5294 genes associated with this cluster, 2,878 are expressed in the early gonad (File 4). Cluster 5 gene regions had no enrichment for H3K27me3 signal, and this was the largest group of genes (17219 genes corresponding to 22163 RefSeq transcripts). Approximately 54% of these genes are expressed at E11.5 (9,447 genes, File 4). When comparing normalized gene expression levels between grouped genes, cluster 3-5 genes are expressed at levels significantly higher than cluster 1 and 2 genes (Fig. 6C, P < 0.0001).

### Genomic regions common to both data sets

To identify regions of enrichment common to both histone marks, peak regions generated by MACSbroadpeakcall for each mark were intersected based on at least 1bp overlap, to produce a list of 12,296 regions (Fig. 7A). Examples of signal distribution for two genes of interest, *Fgf9* and *Esr1*, is shown in Fig. 7C and D, with genome regions common to both marks indicated. These common peak regions were used with GREAT (McLean et al., 2010) to identify genes nearby each genomic region (File 6). These genes may lie in regions of the genome that is being actively transcribed, or either repressed or poised for future activation due to the presence of H3K27me3 in addition to H3K4me3.

**Fig. 7.**
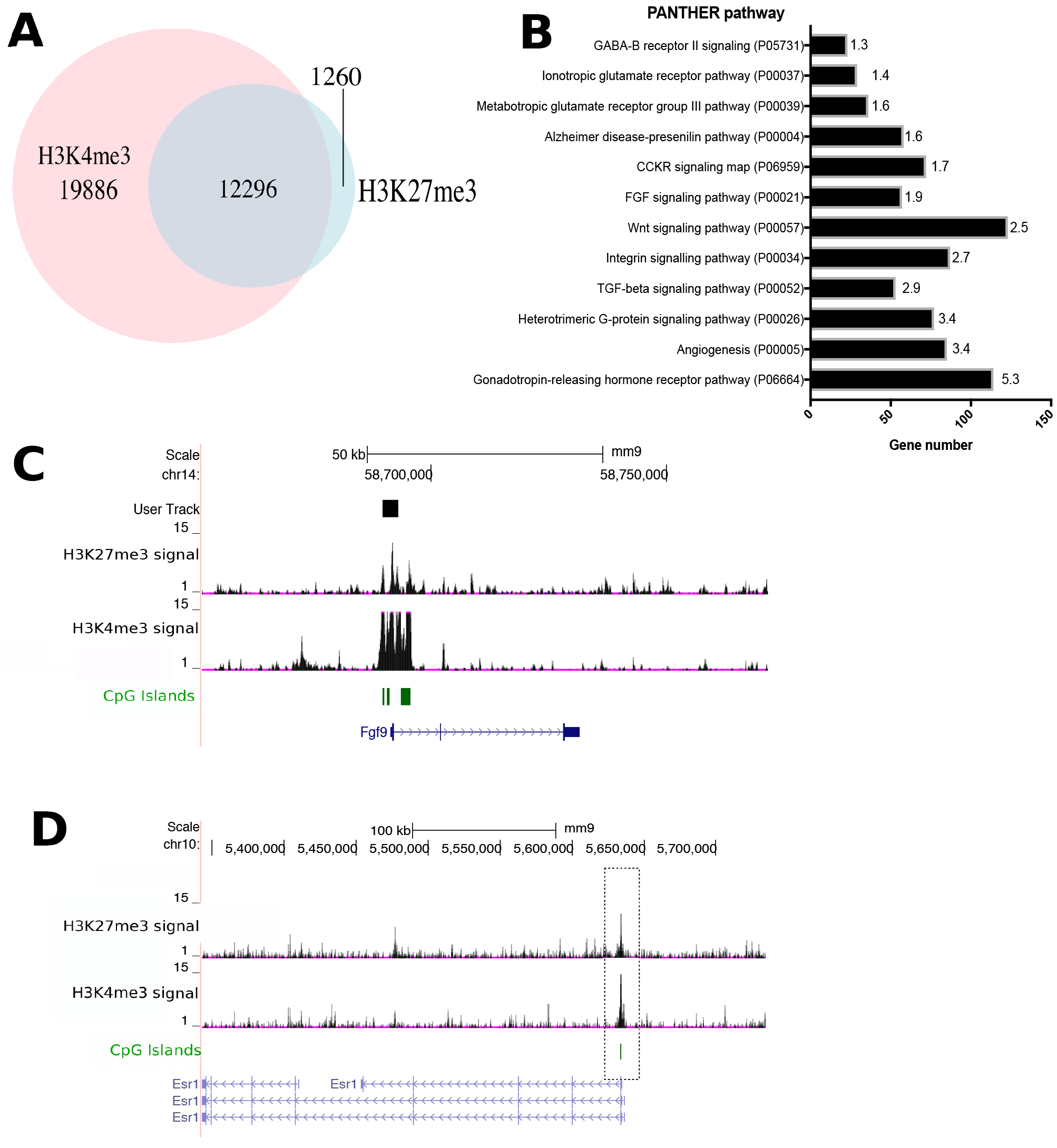
Regions overlap between H3K27me3 and H3K4me3. **A.** Identification of common H3K4me3 and H3K27me3 MACS peak regions. **B.** PANTHER pathway classification for genes associated with H3K4me3-H3K27me3 intersection peak regions. Example of shared peak regions for *Fgf9* (**C**) and *estrogen receptor gene* 1 (*Esr1*, **D**). Regions that intersect between the two peak files are boxed.

The resulting gene list (6523 genes) was used for GO functional annotation in PANTHER (Mi et al., 2010) (Fig. 7B and File 6). PANTHER pathway analysis showed that Wnt, Fgf and Tgfb signaling pathways were significantly overrepresented (Fig. 7B), pathways that have previously been shown to be essential for early gonad development (Colvin et al., 2001; Gustin et al., 2016; Josso and di Clemente, 1999; Kim et al., 2006). In particular, Wnt signaling pathway associated genes (123 genes) were particularly enriched within this list (Fig. 7B). Other pathways of interest included angiogenesis, important for formation of the gonadal blood supply and patterning of the testicular cords during development (Combes et al., 2009; Coveney et al., 2008). Integrin signaling is also required for correct development of the gonad and their primordial germ colonization (Anderson et al., 1999; Messina et al., 2011).

Genes linked to pathways with unknown roles in gonad development were also overrepresented. Gonadotrophin-releasing hormone (GnRH) signaling has been best studied in the adult mouse, where GnRH neuronal signaling peptide loss of function leads to hypogonadism in adult mice. GnRH neuronal signaling does not appear to be functional until after E16.5 (Wen et al., 2010). The *GnRH receptor* (*GnrhR*) gene is expressed in the rat embryonic testis Leydig cells, adult ovary and breast tissues (Ishaq et al., 2013; Schang et al., 2012) but has not been reported to have significant expression in the developing embryo gonad and was included in the array RNA expression data (Jameson et al., 2012). However, the gene for the *Gnrh1* hormone is expressed in developing chick gonad (Carre et al., 2011) and mouse gonad (Fig. S6, (Jameson et al., 2012)).

Several genes linked to glutamate-receptor signaling pathways were also overrepresented on the resulting gene list. These genes are expressed in many non-neural tissues including the adult testis (Julio-Pieper et al., 2011; Marciniak et al., 2016), the developing gonad (Jameson et al., 2012) and are believed to contribute to organ homeostasis. Cholecystokinin receptor (CCKR) pathway components are also overrepresented. While cholecystokinin (a peptide hormone) itself is not expressed in the gonad or has any known function in gonad development, many CCKR downstream pathway genes such as β-catenin, *Jun* and c-Myc are known developmental regulators and are expressed in the developing gonad (Jameson et al., 2012). This may be why this pathway is significantly overrepresented. The CCKR pathway has also been linked to sex dimorphic responses in the brain (Xu et al., 2012).

### DNA sequences associated with H3K4me3 enrichment

To identify candidate factors that might bind to enhancers associated with each histone mark, we used HOMER motif analysis. Narrow peak regions (~1kb) were first identified, as these are more likely to represent a regulatory element rather than global repression/activation of an area of the genome (Fig. 9A). This was followed by HOMER motif enrichment for these narrow peak regions. The HOMER software produces a list of known motifs for transcription factors significantly overrepresented when compared to a set of randomly selected background sequences. This list of enriched motifs was further filtered to those sites whose associated transcription factors are expressed in the developing gonad, by comparison to the available Affymetrix array data (Jameson et al., 2012).

We identified 9332 narrow (1kb) regions with H3K4me3 enrichment (Fig. 8A and File 6). A total of 67 motifs were enriched within these peak sequences, of which 33 of the known motifs correspond to transcription factors that are expressed during gonad development (File 6). Seven of these factors are expressed in a cell-type specific manner (Fig. 8B-H).

**Figure 8.**
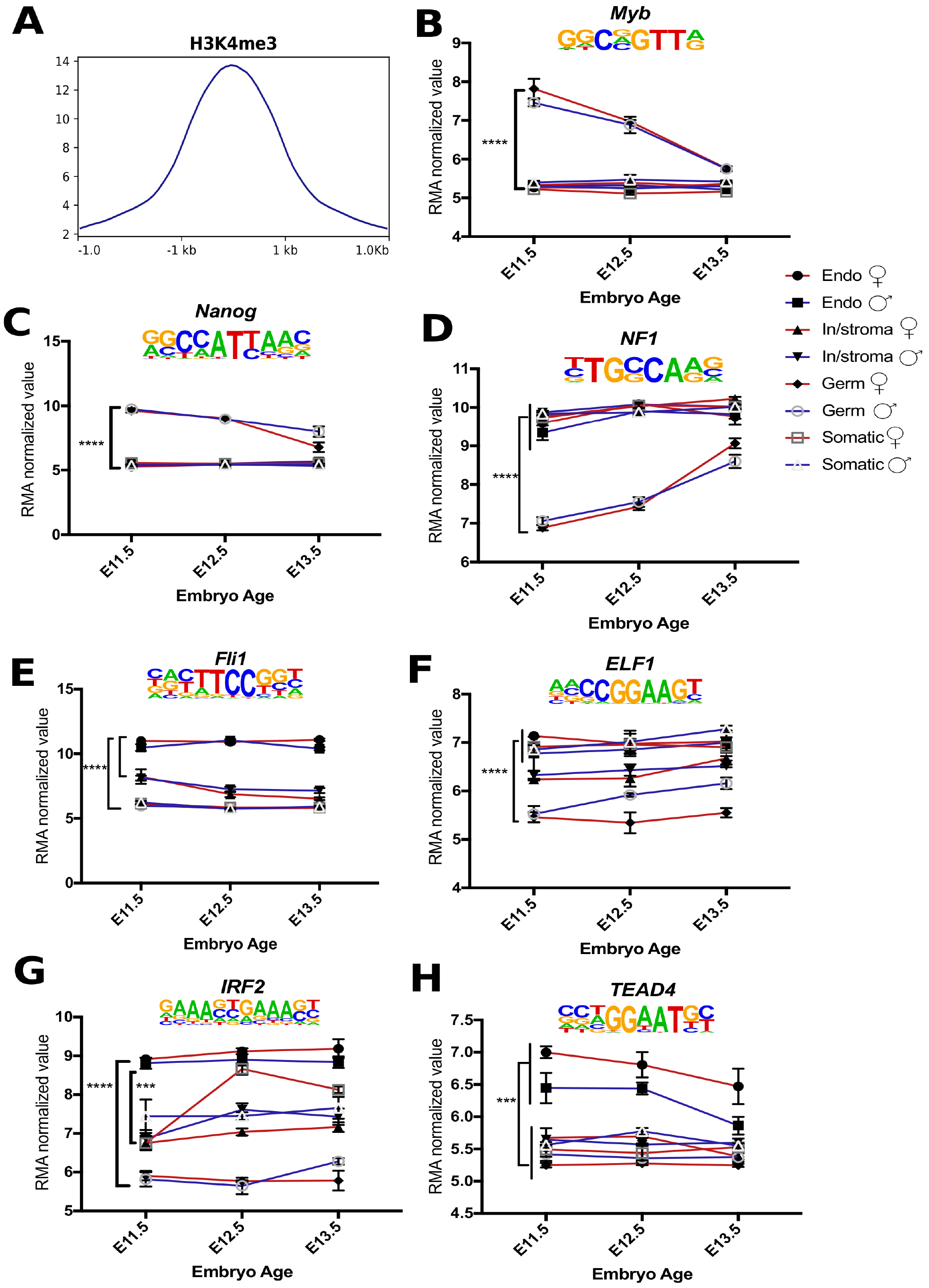
DNA motifs enriched within H3K4me3 HOMER peak regions. **(A**) Signal profile for H3K3me3 across the genome regions (3 kb), centered on the middle of the peak interval. Enriched DNA binding motifs and their cell-lineage expression profiles for germ, interstitial, supporting and endothelial cells at E11.5, E12.5 and E13.5 (Jameson et al., 2012). Profiles are shown for *Myb* (**B**), *Nanog* (**C**), *Nf1* (**D**), *FH1* (**E**), *Elf1* (**F**), *IRF2* (**G**) and *TEAD4* (**H**) genes. Mean +/− standard error of the mean. Statistical analysis: two-way ANOVA Tukey’s multiple comparisons test, adjusted P-value <0.001 (**), <0.0001 (****).

Sixty one percent of H3K4me3 regions had a Myb (c-Myb) DNA binding motif (q-value = 0.007). Myb is a helix-turn-helix transcription factor with roles in cell cycle regulation (Nakata et al., 2007), hematopoiesis (Mukouyama et al., 1999) and oocyte meiotic maturation(Zheng et al., 2012). *Myb* gene expression was higher in GCs at E11.5 compared to other gonadal cell types (Fig. 8B).

Nanog is a well-characterized transcription factor involved in maintaining pluripotency in embryonic stem cells, proliferation and nuclear reprogramming in primordial germ cells (Theunissen et al., 2011; Yamaguchi et al., 2009). Over 80% of H3K4me3peak regions (q-value <0.0001) had potential Nanog binding motifs (File 6). Nanog was expressed at much higher levels in GCs compared to other cell types found in the E11.5 gonad (Fig. 8C).

*Nuclear factor 1* (*NF1*) gene encodes a ubiquitous transcription factor with roles in chromatin remodeling (Gonen et al., 2017) and has been found to function with Sf-1 in adrenal steroidogenesis (Aigueperse et al., 2001). Sixty percent of H3K4me3 peak sequences had a potential NF1 binding site (q-value = 0.02). *Nf1* mRNA is expressed at high levels in all gonadal cell types with the exception of the primordial GCs at E11.5 (Fig. 8D). Expression in male and female GCs eventually increases following sex determination (E12.5) to similar levels of other gonadal cell types by E13.5 (Fig. 8D).

Nine members of the Ets family of transcription factors were identified as binding to motifs located in H3K4me3 narrow peak regions. This protein family has roles in cell lineage specification including blood cell differentiation (Ciau-Uitz et al., 2013; Remy and Baltzinger, 2000). The ETS factor, *Elf1* (*E74-like factor 1*) has an essential role in vascular development (Gaspar et al., 2002; Huang et al., 2006). Around a third (34%) of H3K4me3 narrow peak regions contained *Elf1* binding motifs (q-value <0.0001). *Elf1* mRNA expression levels were lower in germ cells at E11.5, when compared to somatic and interstitial/stroma cells (Fig. 8F, P<0.0001). mRNA for another ETS factor, *Friend leukemia integration 1 (Fli1)*, was expressed at higher levels in endothelial cells (P<0.0001) compared to other cell lineages (Fig. 9E).

**Figure 9.**
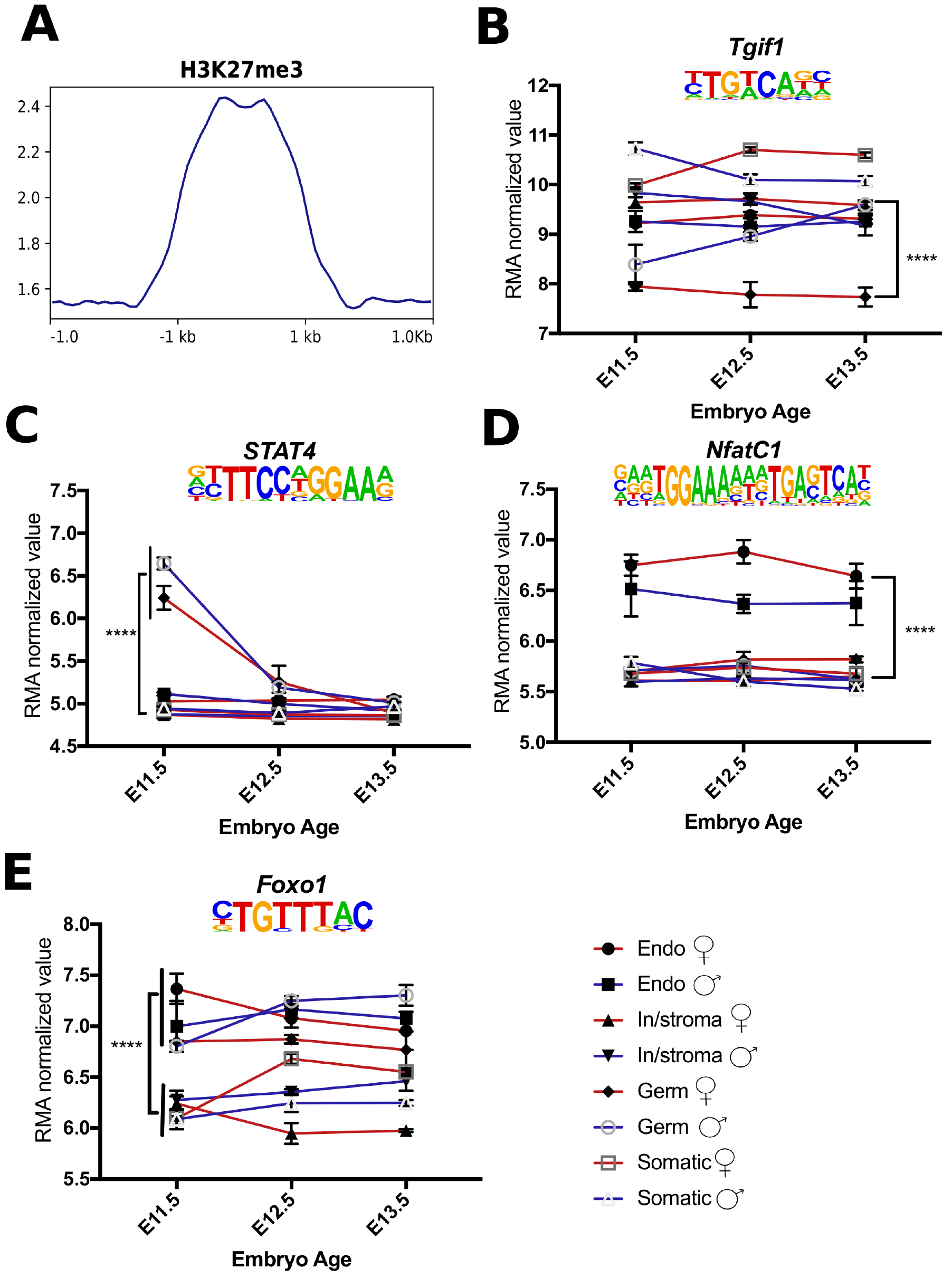
DNA motifs overrepresented within 1kb peak regions enriched with the H3K27me3 modification. **(A)** Signal profile for H3K27me3 across the genome regions (3 kb), centered on the middle of the peak interval. Enriched DNA binding motifs and their cell-lineage normalized expression profiles from E11.5 to E13.5 of gonad development (Jameson et al., 2012). Profiles are shown for *Tgif1* (**B**), *STAT4* (**C**), *NfatC1* (**D**) and *Foxo1* (**E**) genes. Mean +/− standard error of the mean. Statistical analysis: two-way ANOVA Tukey’s multiple comparisons test, adjusted P-value <0.001 (**), <0.0001 (****).

*Interferon regulatory factor 2* (*Irf2*) was expressed at much higher levels in the endothelial cell types, compared to both the somatic and germ cells (Fig. 8G, P<0.001 and P<0.0001 respectively). A small subset of sequences had a Irf2 DNA binding motif (5%, q-value <0.0001, File 6). *Elf-1* and *Irf2* have both been found to induce stem cells to form blood cells in culture (Yamamizu et al., 2013).

The DNA binding motif for TEA domain family member 4 (Tead4, also known as TEF3) was enriched for 30% of sequences (q-value 0.01, File 6). Expression profiling indicates the *Tead4* gene is expressed at higher levels in endothelial cells compared to other cell types present in the gonad (Fig. 8H). Previously, Tead4 was found to directly regulate *Sf1* expression in reporter assays (Sakai et al., 2008), and has been linked to VEGF-mediated angiogenesis (Liu et al., 2011)

### DNA sequences associated with H3K27me3 marks

For H3K27me3, 4,847 regions were used in motif analysis (File 6) and identified enrichment for 15 known motifs for transcriptions factors. Of these, 6 transcription factors are expressed in the E11.5 gonad (Fig. 9B-E).

*TG-interacting factor 1* (*Tgif1*) gene encodes for a homeobox protein that acts as a repressor of TGF-β and RA signaling (Bartholin et al., 2006; Wotton et al., 1999), and can repress embryonic stem cell factors, binding directly to Oct4 (Lee et al., 2015). The majority of H3K27me3 peak regions (88%) had a predicted Tgif1 DNA binding motif (q-value = 0.025). At E11.5, *Tgif1* mRNA is expressed at high levels in the undifferentiated somatic, endothelial and interstitial/stroma tissue, compared to GCs (Fig. 8B, P<0.001). By E13.5, *Tgfif1* mRNA is expressed at higher levels in male GCs, compared to female GCs E13.5 (Fig. 9B, P<0.0001).

*Signal transducer and activator of transcription 4* (*STAT4*) motif was enriched in ~43% (q-value = 0.03) of H3K27me3 narrow peak sequences (File 6). The *STAT4* gene is expressed at higher levels in both male and female germ cells at E11.5, in comparison to other cell types (Fig. 9C, P<0.0001). STAT4 has multiple roles in development, including in endothelial cell proliferation in zebrafish (Meng et al., 2017), and is expressed in the germ cells of the adult mouse testis (Herrada and Wolgemuth, 1997)

The *Nuclear factor of activated T-cells (NfatC1)* gene has roles in lymphatic and valve endothelial development (Kulkarni et al., 2009; Wu et al., 2011). *NfatC1* was expressed at higher levels endothelial cells compared to other cell types in the E11.5 gonad (Fig. 9D, P<0.0001). Nine percent of H3K27me3-associated sequences (q=0.0009) were predicted to contain NfactC1 DNA binding motifs (File 6).

The *Forkhead box protein O1* (*FoxO1*) gene has a role in regulating human and mouse ESC pluripotency through regulation of *SOX2* and *OCT4* gene expression (Zhang et al., 2011), and also functions in the adult mouse male germline and ovarian granulosa cells (Liu et al., 2013). *Foxo1* mRNA levels were higher in GCs and endothelial cells at E11.5 compared to other gonadal cell types (Fig. 9E, P<0.0001). The FoxO1 binding motif was present in 70% of peak sequences (q-value = 0.0001).

## Discussion

We successfully performed ChIP-seq analysis for H3K4me3 and H3K27me3, histone marks that represent two different transcriptional states, on E11.5 UGRs. The resulting signal distribution profiles reflected transcriptional activity in the E11.5 gonad and may indicate areas of the genome either in an active or repressed state, including potential enhancer regions.

Immunofluorescence staining of the UGR revealed that many gonadal cells had higher levels of these histone modifications compared to nearby cells. While some of these cells are likely to be PGCs, others are likely to represent precursors of other cell lineages required for gonad development. Future studies will address how these marks are first established and what happens to these marks over the course of gonadal development. This study used a time-point at the beginning of sex determination and prior to hormone production, and we do not address epigenetic dimorphism between the sexes at E11.5. Therefore, we cannot discount the possibly that sexually dimorphic epigenetic patterns exist prior to sex determination (earlier than E11), and perhaps are based on the karyotype of the embryo. Histone modifications are enzymatically added and removed, therefore the enzymes involved are important regulators of cell differentiation and fate (Butler et al., 2012). While the X and Y-chromosomes both have lysine demethylase genes, known as *Jarid1c* and *Jarid1d* respectively, it is unknown if they have a role in early gonad development. However, in terms of sex-determination, the histone demethylase enzyme, JmjC domain-containing protein (Jmjd1a) was recently shown to be required for correct expression of the *Sry* gene and male sex determination in mice (Kuroki et al., 2017). It will also be important to consider when and how any sex-specific histone modifications are first established during cell-fate specification in the developing gonad (Butler et al., 2012). (Kuroki et al., 2017).

Epigenetic modifications are essential for cell fate and lineage specification. DNA binding motifs located within narrow peak regions included transcription factors that were themselves expressed in a cell-specific manor. This identified several new candidate transcription factors that may have a role in the undifferentiated gonad. Tgif1 is a known repressor of retinoic acid signaling (Bartholin et al., 2006) and is expressed at significantly higher levels in male GCs by E13.5 (Fig. 9B). Repression of RA signaling prevents male GC entry into meiosis (Bowles et al., 2006). Expression of another transcription factor, cMyb was also highest in E11.5 GCs (Fig. 8B). This gene was previously proposed to regulate *KIT* expression (Mithraprabhu and Loveland, 2009), KIT is required for germ cell migration and maturation (Fleischman, 1993).

Another transcription factor gene of interest was *Nfatc1*. Gonad *Nfatc1* gene expression is much higher in endothelial cells (Fig 9D), suggesting it has a role in vascular cell fate in the gonad. Previously, it was shown that *Sox9* inactivation in the mouse heart resulted in ectopic expression of *Nfatc1* in heart mesenchymal cells, and in wild-type animals *Nfatc1* is expressed only in the endothelial cells of the heart (Akiyama et al., 2004). This suggests a negative relationship between *Sox9* and *Nfatc1*, perhaps in restricting cell lineage specific-gene expression.

Several transcription factors predicted to bind to other DNA binding motifs enriched within our dataset were not expressed in the developing gonad based on available array data (File 6). However, these factors may have roles later in development, or in the post-natal gonad. *Recombination signal binding protein for immunoglobulin kappa J region* (*Rbpj*) gene encodes for a Notch pathway regulator, and conditional knockout results in reduced testis size. *Rbpj* is required for the testis stem cell niche (Garcia et al., 2014). Therefore, while *Rbpj* gene expression is detected in the early embryonic gonad, it also affects male fertility later in life.

We determined a set of genomic regions that were enriched with both histone marks in chromatin extracted from E11.5 gonads. Different cell lineages have distinct epigenetic profiles (Hawkins et al., 2010). From this data, we cannot be sure that a given genome region is actually marked by both modifications in the same cell, as we used a mixed population of cells, and we thus are observing the average signal profile over multiple cell types. Therefore, these regions cannot be termed bivalent regions based on this analysis. Future studies, using cell-sorting methods, such as those used by Jameson *et al.*, 2012, and the development of new ChIP methods requiring less material may yield enough sequencing data to examine the epigenetic landscape of individual cell populations within the UGR.

Wnt signaling pathway genes were significantly overrepresented as being epigenetically marked with both modifications. This pathway has roles in formation of the UGR for both sexes, but at later stages of gonad development, it functions in largely in ovarian development only (Boyer et al., 2010; Ottolenghi et al., 2007). Wnt4 and Rpo1 are essential for proliferation of coelomic epithelium in both XY and XX gonads (Chassot et al., 2012). The absence of *Wnt4* in female mice results in masculinization with the development of Wolffian ducts and absence of Müllerian ducts (Bernard and Harley, 2007; Vainio et al., 1999). Further studies have shown that canonical *Wnt4* expression persists in the developing ovary to prevent testicular formation by repressing the SOX9-Fgf9 positive-feedback loop (Kim et al., 2006). *Wnt5a* is required for the development of posterior Müllerian duct structures (the cervix and vagina), primordial germ cell migration and testicular development (Chawengsaksophak et al., 2012; Mericskay et al., 2004).

Overall, this study provides first insight into the epigenetic landscape of the entire UGR at E11.5 and provides an important dataset for further analysis of regulatory elements active during early gonad development

## Methods

### Animals

The University of Otago Animal Ethics Committee granted ethical approval for this study (approvals ET17/12 (November 2012) and ET25/15 (May 2015)). Embryonic development was timed from the presence of a copulation plug, beginning at 0.5 days post coitum (dpc) to account for the period between fertilization and time of plug observation.

### Chromatin immunoprecipitation

Gonad primordium and associated mesonephros tissue was dissected at E11.5 and pooled in 1 mL 1× PBS on ice and fixed in 25 μL 37% formaldehyde to crosslink together the DNA and associated proteins while rocking for 10 min. The crosslinking reaction was stopped with glycine (final concentration of 1 mg/mL). Five litters of crosslinked UGR tissue were pooled together, pelleted by centrifugation and resuspended in 500 μL membrane extraction buffer (20 mM TrisCl pH 8.0, 85mM KCl, 0.5% NP-40) plus 5 μL of proteinase inhibitor (20 μg/μL).

Homogenized tissue was placed on ice for 10 min and centrifuged at 9,000g for 3 min. Pellets were resuspended in 500 μL digestion buffer (50 mM TrisCl pH 8.0, 5mM CaCl_2_, containing 0.5 μL DTT (100 mM) and 1 μL MNase (300 U/μL)) and incubated at 37°C for 30 min, with mixing every 5 min to digest the chromatin. After the incubation, 50 μL of stop solution (200mM EGTA (pH 8.0)) was added and the sample was placed on ice for 5 min. Samples were centrifuged at 9,000g for 3 min and the supernatant removed. To extract the digested chromatin, nuclear extraction buffer (1% SDS, 10 mM EDTA pH 8.0, 50 mM TrisCl pH 8.0) was added to the pellet to give a final volume of 250 μL and sample left on ice for 15 min, with vortexing every 5 min for 15 s. Samples were centrifuged at 9,000g for 3 min, with the supernatant now containing the desired fragmented chromatin. 10 μl was retained to serve as the “input” control sample for sequencing.

To bind the antibodies to the magnetic beads, 25 μL of Dynabeads (Life Technologies) were washed 3 times in block solution (0.5% BSA in PBS). 10 μg of primary antibody (either anti-H3K27me3 or anti-H3K4me3) in 1 mL block solution was added to the Dynabeads. Mock tubes were set up without primary antibodies, containing 1 mL of block solution, 25 μL Dynabeads and preimmune serum. Beads were rotated at 4°C overnight, washed 3 times in block solution to remove unbound antibody and resuspended in 1.25 mL block solution. Equal volumes of the chromatin-containing supernatant was added to the primary antibody or mock tubes. Microfuge tubes containing Dynabeads were left to rotate at 40C overnight. Antibodies used for ChIP and immunofluorescence were anti-H3K4me3 (ab8580) and anti-H3K27me3 (ab6002), both from Abcam.

Following immunoprecipitation, the beads were washed five times with wash buffer 1 (50 mM HEPES-KOH (pH 7.5), 500mM LiCl, 1 mM EDTA, 1.0% NP-40 and 0.7% sodium-deoxycholate), followed by two washes with wash buffer 2 (TE with 50 mM NaCl). Bound nucleoprotein complexes were extracted with elution buffer (50 MM TrisCl pH 8.0, 1% SDS) at 65°C for 10 min. Protein-DNA cross-linking was reversed for input and ChIP fragments and proteins digested by Proteinase-K (1 μg) 65°C for 90 min, followed by phenol-chloroform extraction. DNA samples were used to construct Thruplex DNA libraries and sequenced on a Illuminia HiSeq platform generating 50 bp single end reads. Sequencing reads were aligned to the mouse genome (mm9) prior to peak calling.

### ChIP-seq data analysis

Sequences returned were assessed for quality by Phred(Q) score. High quality reads were aligned to the mouse genome (mm9) using Map with BWA (Galaxy Version 0.3.1, with default settings). Poorly mapped reads (MAPQ score <20) were removed by filtering with samtool_filter2 (1.1.1). To examine correlation between replicates, a correlation matrix was generated using multiBamSummary (2.4.1) with a bin size of 10000. The correlation heatmaps were plotted using plotCorrelation (2.4.1), using Pearson correlation.

To generate BigWig files for signal visualization and bedgraph files for MACS2 analysis, ChIP-seq output (plus antibody) aligned reads were normalized to the Input signal using bamCompare (2.5.0) using the following settings: 50 bin, readCount set to scale samples to the smallest dataset and ratio to compare to input sample.

To annotate signal with respect to gene features, HOMER and CEAS software packages were used (Heinz et al., 2010)(Heinz et al., 2010; Shin et al., 2009). HOMER was used to annotate genes and gene regions (annotatepeaks function, tag count distribution across the gene body). CEAS (1.0.0) analysis was carried out using default settings (bin size = 50) with previously generated bigwig files and MACSbroadcall peak regions for each histone mark.

The Deeptools (Ramirez et al., 2014) package was firstly used to generate a matrix (computematrix (2.5.0)) to calculate the signal around the gene reference point (start of the region, TSS as the reference-point with a bin size of 50). PlotProfile was used to plot the signal over gene regions. Heatmaps were created using plotHeatmap (2.5.0, averageTypeSummaryPlot=mean, with kmeans clustering).

MACS2 (Feng et al., 2012) bdgbroadcall package (2.1.1) using the normalized (to input) bedgraph file to call variable size peak regions, with the following settings: cutoff for peaks set as 5, minimum length of peak to 200, maximum gap between significant peaks was set to tag size (50) and maximum linking between significant peaks of 1500.

### RNA expression data analysis

Robust Multi-array Average (RMA) normalized values for each gene and cell lineage were obtained from Jameson et al., supplementary data files (Jameson et al., 2012). This study used mouse lines with cell-specific markers and fluorescence-activated cell sorting (FACS) to isolate difference cell populations. Markers used were: *Sry*-EGFP/*Sox9*-ECFP supporting cells, *Mafb-EGFP* to isolate interstitial (XY) and stromal cells (XX), germ cells were isolated using *Oct4*-EGFP and endothelial cells with *Flk1*-mCherry. Gene expression were analyzed using a Affymetrix Genechip 1.0 ST array (Jameson et al., 2012) and reported as normalized RMA values.

### Motif identification using HOMER

HOMER (Heinz et al., 2010) was used to create tag libraries for each ChIP-seq data set. Findpeaks (part of the HOMER package) was used to find genomic regions of ~1kb signal enrichment compared to surround regions, with following parameters: -minDist 2500 - tagThreshold 25 -L 0. The coordinates of the regions identified for histone marks are listed in additional data File 6. To find motifs within HOMER 1kb ChIP-seq regions, we used findMotifs (HOMER) using default parameters (-size given).

### PANTHER GO and Pathway analysis

The RefSeq transcript list for each signal profile cluster was used for PANTHER GO-Slim-Biological processes with Fisher’s Exact with FDR multiple test correction. The returned GO annotations and their adjusted p-values were visualized using Revigo and Treemap (Supek et al., 2011).

Genomic coordinates for regions that enriched within both MACS peak data sets were submitted Genomic Regions Enrichment of Annotations Tool (GREAT (McLean et al., 2010)) using default parameters and mouse genome (mm9) as background, to generate a list of genes nearby and overlapping with each peak region. This gene list was inputted into PANTHER for GO analysis (File 5). PANTHER pathway analysis P-values are adjusted for Bonferroni correction.

### Immunofluorescence of paraffin sections

Sections of E11.5 embryos were de-paraffinized and rehydrated as described previously (Yang and Wilson, 2015). For antigen retrieval, slides were places in sodium citrate buffer (pH 6.0) and heated in a microwave for 30 min. Slides were left to cool at room temperature and then rinsed in dH_2_O. Slides were washed three times with PBTx (PBS with 0.1% Triton X-100) and then blocked with 10% heat inactivated sheep serum in PBTx for 2 h at room temperature. Primary antibodies were diluted in blocking solution and incubated overnight at 4°C. For each secondary, a control staining (no primary antibody) was performed. Samples were washed with PBTx three times to remove unbound primary antibodies and then blocked for 20 min. Secondary antibodies were diluted in blocking solution and added to the slides before incubation at room temperature for 2 h. Samples were washed three times with PBTx before mounting. Staining was imaged on a Zeiss LSM 710 confocal microscope.

Secondary antibodies used were donkey anti-rabbit (Alexa Fluor 488, Life Technologies) and anti-mouse (DyLight^®^ 488, Abcam) to detect anti-H3K4me3 and anti-H3K27me3 respectively. Antibody dilutions used were as follows: anti-H3K4me3 (1 in 50), anti-H3K27me3 (1 in 50), donkey anti-rabbit (1 in 100), donkey anti-mouse (1 in 400).

## Data access

ChIP-seq data has been deposited in Gene Expression Omnibus (GEO) with the accession number GSE109380.

## Acknowledgements

Research described here was funded by a University of Otago Research Grant to Megan Wilson. Yisheng Yang was supported by a University of Otago PhD scholarship. We thank James Smith, Susie Szakats, Kathy Sircombe, Jeremy McCallum-Loudaec, Rebecca Clarke and Lyvianne Decouryte for feedback on manuscript drafts.

## Supplementary Data files

**File 1. Sequencing data mapping statistics.**

**File 2. List of regions with similar signal distribution identified by deepTools/plotHeatmap using k-means clustering analysis.** Output file giving region coordinates, nearby feature/gene and cluster group. Sheet1: H3K27me3 signal clusters. Sheet 2: H3K4me3 signal clusters.

**File 3. PANTHER Gene ontology analysis for H3K4me3 clusters.** Each sheet summarizes the PANTHER analysis carried out for the cluster gene list. Each gene list includes genes associated with each signal profile (cluster).

**File 4. PANTHER Gene ontology analysis for H3K27me3 clusters.** Each sheet summarizes the PANTHER analysis carried out for the cluster gene list. Each gene list includes genes associated with each signal profile (cluster).

**File 5. MACS broadcall peak regions.** Excel file with the peak regions identified for each histone mark and a list of peak regions that are shared between both marks (“overlapping regions”). Sheet 1: MACS broadcall peaks for H3K4me3. Sheet2: MACS broadcall peak regions for H3K27me3. Sheet 3: Overlapping peak regions. Sheet 4: DAVID annotation analysis (summarized in Fig. 7B). Sheet 5: Genes located near shared peak regions (identified using GREAT).

**File 6. Motif analysis results for narrow peak regions.** 1 kbp peak regions identified by HOMER analysis for H3K27me3 and H3K4me3. Motifs Homer analysis for known DNA binding motifs. Grouped into motifs of factors expressed in the early gonad (Jameson et al., 2012) and those factors not present in array data by Jameson *et al.* 2012.

Supplementary figures are provided in a single pdf file.

